# Hand2 represses non-cardiac cell fates through chromatin remodeling at *cis-*regulatory elements

**DOI:** 10.1101/2023.09.23.559156

**Authors:** Valerie Komatsu, Brendon Cooper, Paul Yim, Kira Chan, Wesley Gong, Lucy Wheatley, Remo Rohs, Scott E. Fraser, Le A. Trinh

**Author notes:** Centre for Clinical Brain Sciences, Chancellor’s Building, University of Edinburgh EH16 4SB (Lucy Wheatley, Scott E. Fraser).

## Abstract

Developmental studies have revealed the importance of the transcription factor Hand2 in cardiac development. Hand2 promotes cardiac progenitor differentiation and epithelial maturation, while repressing other tissue types. The mechanisms underlying the promotion of cardiac fates are far better understood than those underlying the repression of alternative fates. Here, we assess Hand2-dependent changes in gene expression and chromatin remodeling in cardiac progenitors of zebrafish embryos. Cell-type specific transcriptome analysis shows a dual function for Hand2 in activation of cardiac differentiation genes and repression of pronephric pathways. We identify functional *cis-*regulatory elements whose chromatin accessibility are increased in *hand2* mutant cells. These regulatory elements associate with non-cardiac gene expression, and drive reporter gene expression in tissues associated with Hand2-repressed genes. We find that functional Hand2 is sufficient to reduce non-cardiac reporter expression in cardiac lineages. Taken together, our data support a model of Hand2-dependent coordination of transcriptional programs, not only through transcriptional activation of cardiac and epithelial maturation genes, but also through repressive chromatin remodeling at the DNA regulatory elements of non-cardiac genes.

## Introduction

The developing heart has provided a powerful and accessible system for understanding how differentiated cell types are specified during embryonic development. Cardiogenesis begins with the specification of cardiac progenitor cells (CPCs) within the lateral plate mesoderm (LPM), an embryonic tissue that gives rise to the heart, blood vessels, and blood cells (Stainier, Lee et al. 1993, Schoenebeck, Keegan et al. 2007, Mao, Boyle Anderson et al. 2021). The LPM sits as paired primordia on the left and right sides of the embryo, bordered by non-cardiac tissues: the intermediate mesoderm (IM), ectoderm and endoderm, tissues that form urogenital structures, epithelial and neural tissues, and digestive and respiratory tubes (Mallo, Wellik et al. 2010). These neighboring tissues play important roles in signaling to the LPM to define the cardiac field. The endoderm secretes bone morphogenic protein (BMP) and fibroblast growth factor (FGF), which pattern the LPM, creating a permissive environment for myocardial gene expression (Komuro and Izumo 1993, Schoenebeck, Keegan et al. 2007). The notochord and ectoderm secrete inhibitory signals to limit the expansion of the cardiac fields (Schultheiss, Burch et al. 1997, Alsan and Schultheiss 2002). These permissive and inhibitory signals promote and restrict the cardiac tissue field early in development, helping to define the subregion of the LPM with the potential to become CPCs.

Although undifferentiated, CPCs are committed to a cardiac cell fate early in development. Through the expression of transcription factors such as Nkx2.5, Tbx20, Gata5, and Hand2, CPCs activate gene networks that turn on expression of myocardial structural genes, including myosin light and heavy chains, cardiac troponins, tropomyosin, and titin, resulting in the differentiation of CPCs into cardiomyocytes (Reiter, Alexander et al. 1999, Yelon, Ticho et al. 2000, Stennard, Costa et al. 2005, Holtzinger and Evans 2007, Guner-Ataman, Paffett-Lugassy et al. 2013, Lu, Langenbacher et al. 2017). In zebrafish, CPC differentiation begins at the 10-somite stage (10ss; 14 hours post fertilization, hpf), at the onset of LPM migration towards the embryonic midline, where the heart tube will later form (Bakkers 2011, Lou, Deshwar et al. 2011, Guner-Ataman, Paffett-Lugassy et al. 2013). During LPM migration, CPCs undergo a mesenchymal-to-epithelial transition (MET) (Orts-Llorca 1963, Lough and Sugi 2000). This epithelial maturation couples the migration and differentiation of CPCs, and plays a central role in heart morphogenesis, as it is required to form the myocardial epithelium and the linear heart tube (Trinh and Stainier 2004).

Hand2, a basic helix-loop-helix (bHLH) transcription factor, is critical for both CPC differentiation and epithelial maturation, in addition to playing key roles in branchial arch and limb bud development (Yelon, Ticho et al. 2000, Trinh, Yelon et al. 2005). Hand2 has been studied largely in the context of cardiac differentiation, documenting its role as a transcriptional activator (Srivastava, Thomas et al. 1997, Yelon, Ticho et al. 2000, Dai, Cserjesi et al. 2002, Trinh, Yelon et al. 2005, Holler, Hendershot et al. 2010, Laurent, Girdziusaite et al. 2017). Hand2 has also been reported to have repressive roles during development; for example, during atrioventricular canal (AVC) formation in mouse, Hand2 directly represses a number of genes involved in the epithelial-to-mesenchymal transition (EMT) of AVC cells (Laurent, Girdziusaite et al. 2017). Other studies in developing mice show that Hand2 mediates the repression of *Dlx5* and *Dlx6* in mandibular arches that is essential for proximal-distal patterning of the lower jaw of *Tbx18* and *Irx3/5*, refining the proximal boundary of the limb bud mesenchyme (Barron, Woods et al. 2011), and of pronephros differentiation through restricting the IM in zebrafish and avians (Perens, Garavito-Aguilar et al. 2016).

Despite the clear evidence for Hand2 as a key transcriptional regulator, few studies have assessed Hand2 function at *cis*-regulatory elements (CRE) in CPCs. Potential tissue-specific enhancers have been identified by ATAC-seq (Assay for Transposase Accessible Chromatin sequencing) (Daugherty, Yeo et al. 2017, Uyehara, Nystrom et al. 2017, Yuan, Song et al. 2018). Chromatin accessibility assays during fibroblast reprogramming into cardiomyocyte-like cells have revealed Hand2-dependent chromatin reorganization, suggesting that Hand2 regulation of target genes involves epigenetic remodeling (Fernandez-Perez, Sathe et al. 2019). Chromatin accessibility assays at the onset of CPC differentiation have identified potential LPM-specific CREs (Yuan, Song et al. 2018); however, parallel studies of the changes in the transcriptome or epigenome after the depletion of Hand2 have not been performed. Therefore, the relative roles of Hand2-regulated chromatin remodeling and Hand2 interactions with *cis* regulatory elements remain unexplored during CPC differentiation and epithelial maturation.

Here, we address these important open questions by assessing Hand2-dependent transcriptomic and epigenomic changes within the LPM by comparing normal LPM cells to those with CRISPR-induced mutations within *hand2* (*hand2* crispants). Our data reveal that Hand2 promotes not only transcriptional activation of genes involved in CPC differentiation and epithelial maturation, but also represses non-cardiac associated gene expression through chromatin remodeling of their enhancers. These findings validate Hand2’s role in cardiogenesis through activating genes involved in CPC differentiation and epithelial maturation, in addition to expanding its role to include transcriptional repression in the LPM through chromatin remodeling. This suggests a novel mechanism by which Hand2 ensures proper cardiac development by simultaneously repressing non-cardiac enhancers and activating transcription of cardiac differentiation and epithelial maturation genes, revealing a robust embryonic mechanism for defining and reinforcing tissue fields in early development.

## Results

### Specific labeling of myocardial progenitor cells with hand2 enhancer driven transgene

To enable the isolation of CPCs at the onset of cardiac differentiation, we created a transgene that expresses eGFP under the control of a *hand2* enhancer element and minimal *e1b* promoter, *Tg(hand2enh:eGFP)* (Fig. 1A). This transgene labels the anterior and posterior LPM at 10-somites (14 hpf), the stage when CPCs are specified within the anterior LPM (Fig. 1B). Expression of the eGFP reporter persists in myocardial cells as the heart tube forms and develops (Fig. 1), permitting CPC identification and isolation throughout early stages of heart development.

**Fig. 1:**
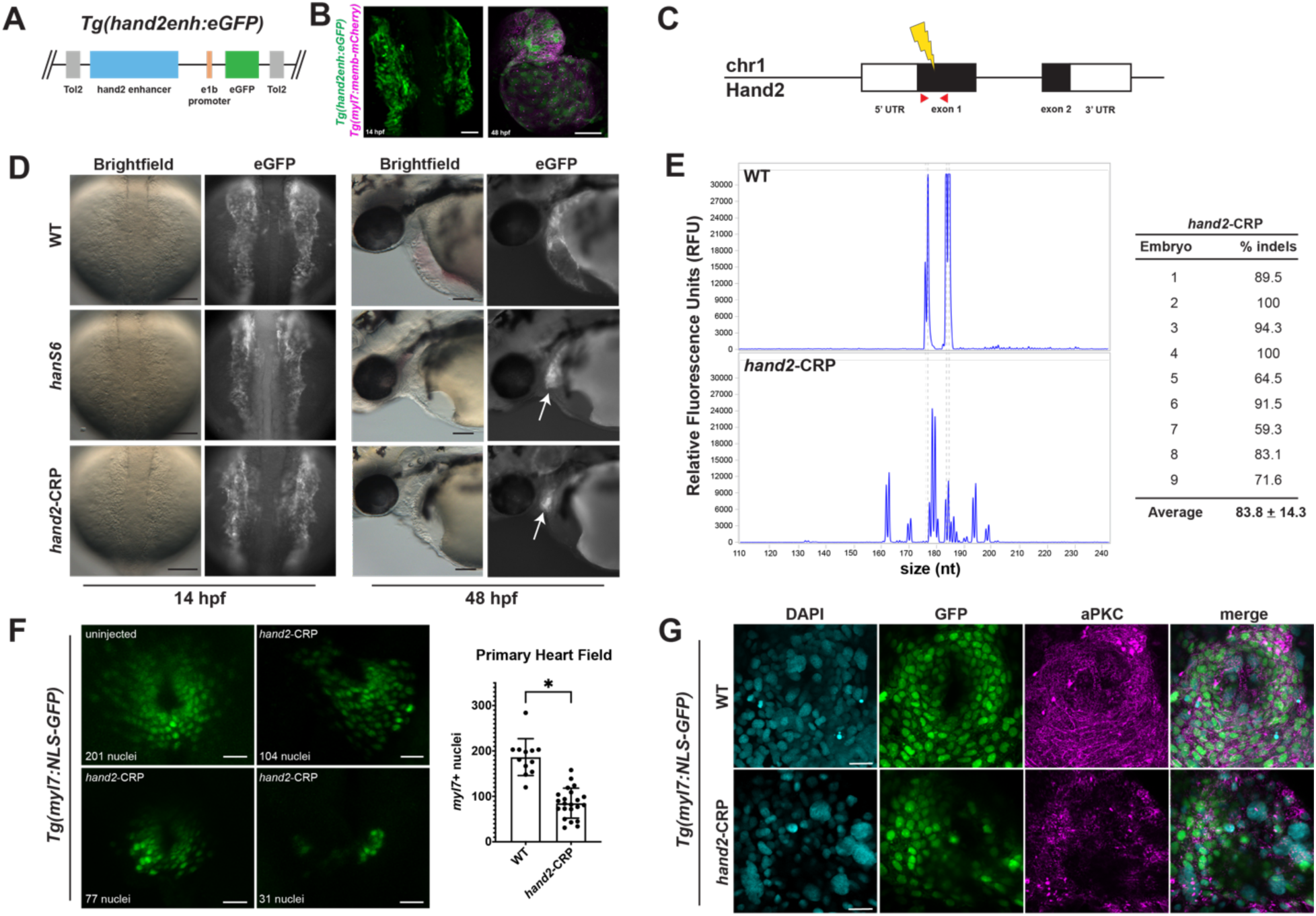
CRISPR-Cas9 induced mutagenesis recapitulates homozygous null *Hand2* mutant phenotype. (A) Schematic representation of the *hand2enh* reporter construct. Construct contains a putative enhancer 28Kb upstream of the *hand2* TSS (blue rectangle; *hand2enh*), an *e1b* minimal promoter (orange) and eGFP (green), flanked by minimal *Tol2 cis* sequences to permit integration by Tol2 transposase. (B) Confocal images of reporter expression in the LPM of a 14hpf *Tg(hand2enh:eGFP*) embryo (dorsal view, anterior to the top), and in the heart of a 48 hpf *Tg(hand2enh:eGFP)*; *Tg(myl7:cherry-Ras)* embryo (ventral view). (C) Depiction of the *Danio rerio hand2* locus with the CRISPR-Cas9 target site used to generate *hand2* crispants at 260bp into exon 1, marked with a yellow lightning bolt. PCR primers used for fragment analysis to confirm CRISPR-induced mutagenesis are shown as red arrowheads. (D) Epifluorescence images of *Tg(hand2enh:eGFP)* embryos at 14 (dorsal view, anterior to the top) and 48 hpf (lateral view, anterior to the left): wildtype (WT), *hand2* mutant (*hanS6*), and *hand2* crispant (*hand2*-CRP). White arrows indicate residue myocardial tissue in *hand2* mutant and crispant. (E) Representative capillary electrophoresis traces of WT and *hand2*-CRP embryos, plotted as the number of bases from the start of the amplified fragment. The percentage of cells with indel mutations for 9 representative embryos is indicated at right. (F) Confocal images of control (uninjected) and crispant *Tg(myl7:nucGFP)* embryos at 20-ss (19 hpf), ventral view. *hand2* crispants have significantly fewer myocardial cells than wildtype embryos (unpaired two-tailed t-test; *p < 0.0001). (G) Confocal images, ventral view, of whole mount *Tg(myl7:nucGFP)* embryos at 19 hpf. WT and *hand2* crispant transgenic embryos, immunostained with an antibody against aPKCs (purple) and nuclear-counterstained with DAPI (cyan); myocardial cells are labeled by myl7-driven nuclear *GFP* expression (green). Scale bars: 50um in B, D; 30um in F; 20um in G.

### *Hand2* CRISPR-Cas9 induced F_0_ mutants

To identify *hand2*-dependent genes, we developed an experimental pipeline for differential genomic profiling, comparing wildtype and Hand2-deficient LPM cells. Genomic analysis of Hand2-deficient CPCs is challenging, as null *hand2* mutants (*hanS6*) are embryonic lethal, requiring that stocks be maintained as heterozygotes, limiting the efficiency of null embryo recovery to 25% (Yelon, Ticho et al. 2000). More importantly, during the stages of cardiac progenitor specification, homozygous null (*han^s6/s6^)* embryos are morphologically indistinguishable from their heterozygous (*han^s6/+^*) and wildtype siblings *(han^+/+^*; Fig. S1) (Yelon, Ticho et al. 2000) (Fig. 1D). The absence of a robust morphological phenotype at these early stages makes it challenging to isolate *han^s6/s6^*embryos for genomic profiling.

To obtain Hand2-deficient embryos throughout CPC development with high-confidence, we employed an efficient F_0_ CRISPR-Cas9 mutagenesis system (Jacobi, Rettig et al. 2017, Hoshijima, Jurynec et al. 2019), using chemically modified crRNAs and tracrRNAs, preloaded in Cas9 protein, to create a dgRNP complex for injection into embryos. The chemically modified crRNAs and tracrRNAs resist nuclease-driven degradation (Hendel, Bak et al. 2015, Schubert, Cedrone et al. 2018). Injection of dgRNP complexes with crRNAs targeting *hand2* exon 1 (Fig. 1C) into *Tg(hand2enh:eGFP*) embryos resulted in morphologically-normal 14 hpf embryos (*hand2*-CRP, Fig. 1D); however, when scored at 48 hpf, the *hand2*-CRP embryos exhibited dramatic cardiac defects, including a loss of myocardial cells (Fig. 1d), strikingly similar to homozygous null (*han^s6/s6^*) embryos. Heart defects were observed in over 90% of *hand2*-CRP embryos, similar to the rates reported for a previously validated Hand2-translation blocking morpholino (Maves, Tyler et al. 2009) (Fig. S2). Fragment length analysis by capillary electrophoresis on individual *hand2*-CRP embryos validated the efficiency of mutagenesis at the *hand2* locus following *hand2* dgRNP injection: over 83% of the cells in individual *hand2* F_0_ CRISPR embryos contained indels at the crRNA target site (Fig. 1E; Fig. S3). This high rate of Cas9-induced modification at the *hand2* locus, and the high phenotypic penetrance of cardiac defects, in *hand2*-CRP embryos suggest that injection of *hand2*-targeting dgRNP complexes are an efficient means to obtain homozygous null mutant embryos.

To further evaluate the phenotypic differences between wildtype, *hand2*-CRP and *hand2* null embryos at stages of CPC development, we assessed myocardial cell numbers and epithelial polarity in transgenic embryos expressing nuclear-localized *GFP* in differentiated myocardial cells (*Tg(myl7:nucGFP);* Lu, Langenbacher, & Chen, 2017). The *hand2*-CRP embryos had significantly fewer myocardial cells than their wildtype siblings (Fig. 1F), similar to the dramatic reduction in the number of cardiac progenitors and the loss of apical-basal polarity of the myocardial epithelium in *hand2* null mutants (Trinh, Yelon et al. 2005). Interestingly, we observed three different types of myocardial cell loss in *hand2*-CRP embryos. The first class had approximately half the number of myocardial cells as wildtype, exhibiting a loss of either the left or right side of the heart field (Fig. 1F, top right panel). The second class lacked chamber-specific precursors, predominantly an absence of the lateral atrial precursors (Fig. 1F, bottom left panel). The third and most severe class displayed a dramatic reduction in both left and right precursors, including ventricular precursors (Fig. 1F, bottom right panel). Such variations might reflect the rate of Cas9-induced mutagenesis and perdurance of residual functional Hand2 in the F_0_ embryos. To assess whether any remaining myocardial cells in *hand2*-CRP embryos are able to form a polarized epithelium, we performed immunostaining for the polarity protein, atypical protein kinase C (aPKCs) in *Tg(myl7:nucGFP)* transgenic embryos. We observed a loss of aPKC’s localization to the junctional contacts between myocardial cells in *hand2*-CRP embryos, as previously shown in *hanS6* embryos (Fig. 1G) (Trinh, Yelon et al. 2005), suggesting that the remaining myocardial cells have reduced Hand2 function. Taken together, these data demonstrate that injection of *hand2* dgRNP complexes induces canonical *hand2* mutant phenotypes with high efficiency.

### Gene expression analyses reveal Hand2 transcriptional repressor and activator functions

To identify Hand2-dependent genes during CPC specification and epithelial maturation, we performed RNA-seq on LPM cells isolated from wildtype and *hand2*-CRP *Tg(hand2enh:eGFP)* embryos using fluorescence activated cell-sorting (FACS; Fig. 2A). Despite a reduction in myocardial cells in *hand2*-CRP embryos (Fig. 1F), wildtype and *hand2*-CRP embryos had similar numbers of LPM *eGFP+* cells, indicating that the *Tg(hand2enh:eGFP)* reporter expression is independent of functional Hand2 (Fig. S4). Chromatin accessibility analysis of the genomic *hand2* enhancer element used to generate the transgene shows no changes in ATAC peaks in either wildtype or *hand2*-CRP datasets, demonstrating that chromatin accessibility at this locus does not require Hand2 function (Fig. S6). These data suggest that this *hand2* enhancer is upstream of Hand2 and unaffected by reduced Hand2 function.

**Fig. 2:**
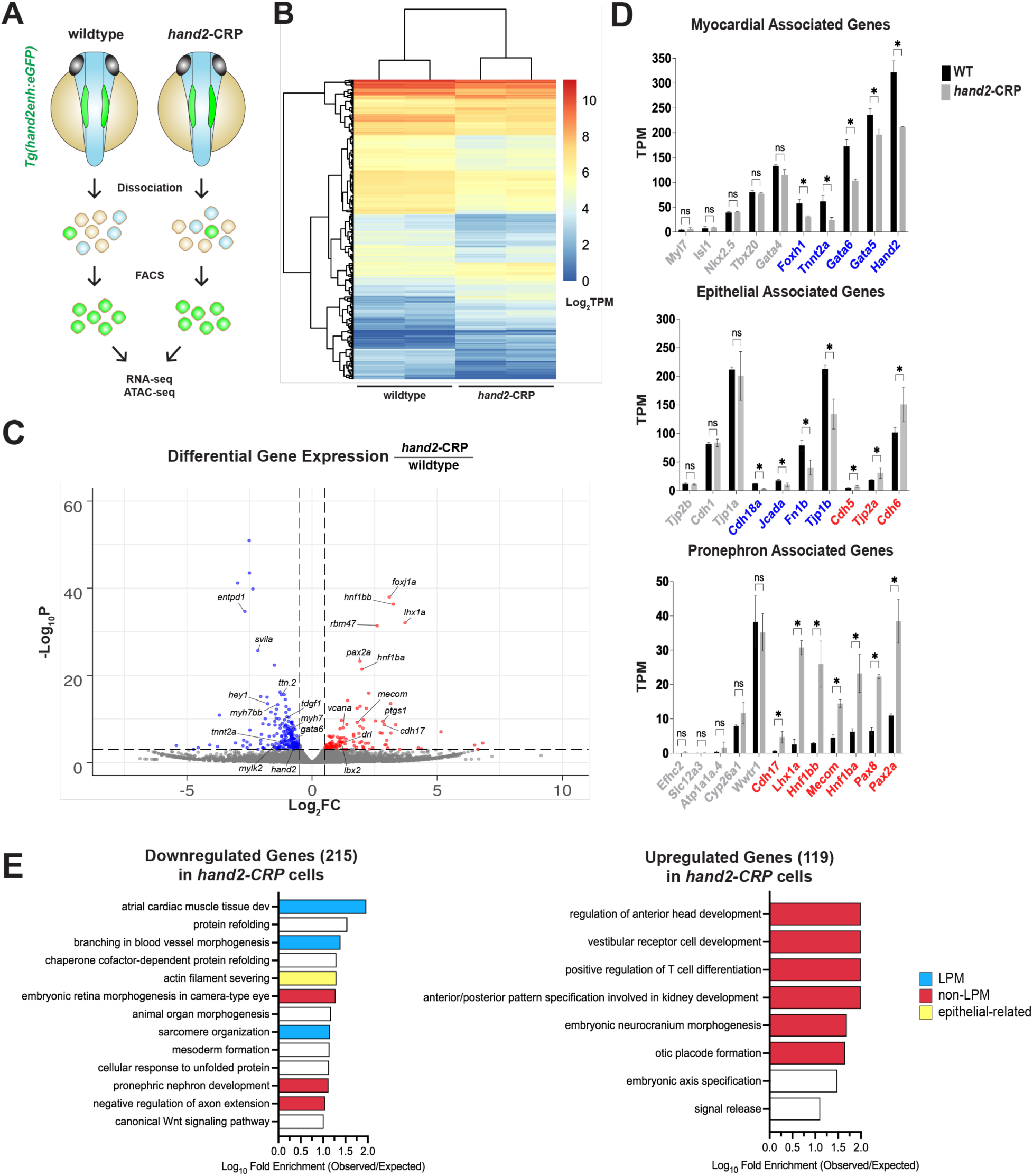
Tissue-specific enrichment of gene expression in *hand2* mutant LPM cells. (A) Schematic illustrating experimental workflow for differential RNA-seq and ATAC-seq analysis of wildtype vs *hand2*-CRP LPM cells. Control (uninjected) and *hand2* crispant (*hand2*-CRP) *Tg(hand2enh:eGFP)* embryos were dissociated and FACS-sorted for *eGFP*. eGFP positive isolated cells were split for either RNA-seq or ATAC-seq library preparations. (B) Heatmap of differentially expressed genes (DEGs) between control and *hand2* crispants with hierarchical clustering. Log2-fold enrichment color key shown to the right of the heatmap. (C) Volcano plot depicting the relative expression of significantly differentially expressed genes in *hand2*-CRP compared to wildtype LPM cells. Significant DEGs (defined as p ≤ 0.001 and log_2_FC ≥ |0.5|) are indicated by dotted lines, red (enriched in *hand2-CRP*) or blue (depleted in wildtype) dots. Nonsignificant DEGs are indicated by gray dots. (D) TPM levels of genes implicated in three distinct differentially expressed gene networks, associated with the myocardium, epithelium or pronephros. Significantly differentially expressed genes (*, p-adj < 0.05) are indicated in blue or red if they are depleted or enriched, respectively, in *hand2*-CRP LPM cells compared to wildtype. Non-significantly differentially expressed genes (ns = not significant) are indicated in gray. (E) Gene ontology (GO) enrichment analysis of biological processes for downregulated (215 genes) and upregulated (119 genes) DEGs in *hand2-CRP* LPM cells. GO subclasses with a Log_10_ fold enrichment > |1.0| were classified as either LPM (process occurring only in LPM, cyan), non-LPM (process excluded from the LPM, red), or epithelial-related (process involved in regulation of epithelial cell polarity, yellow). General biological processes found in both LPM and non-LPM tissues are indicated in white.

Differential expression analysis of wildtype and *hand2*-CRP LPM cell RNA-seq datasets identified 334 differentially expressed genes (DEGs) (p<0.001): 215 genes were significantly depleted in *hand2*-CRP datasets; 119 genes were significantly enriched (Fig. 2B, Table S3). Cardiac genes, including *myh7, myh7bb, mylk2, ttn.2,* and *tnnt2a,* were significantly depleted in *hand2*-CRP LPM cells, consistent with Hand2’s role as an activator of myocardial differentiation (Fig. 2C,D, Table S3). *hand2* itself was slightly downregulated (-0.484 log_10_FC) in *hand2*-CRP datasets, suggesting possible autoregulation as observed during limb bud development (Osterwalder, Speziale et al. 2014). Among those significantly enriched in the *hand2*-CRP datasets were genes marking subregions of the LPM such as *drl, foxj1a, vcana, lbx2,* and *ptgs1* (Fig. 2C, Table S3). Additionally, genes implicated in pronephros differentiation, such as *lhx1a, pax2a,* and *cdh17* were enriched in the absence of Hand2 (Fig. 2C), consistent with a previously described role for Hand2 in the restriction of IM formation (Perens, Garavito-Aguilar et al. 2016).

To identify molecular pathways potentially modulated by Hand2, we explored the differential expression of genes associated with myocardial, pronephros or epithelial differentiation (Fig. 2D). Many of the evolutionarily conserved transcription factors critical for early heart development (including *foxh1, gata5,* and *gata6)* were depleted in *hand2*-CRP datasets (Fig. 2D, top plot). Conversely, *mecom, hnf1bb, pax8*, and other key regulators of pronephros differentiation were enriched in the *hand2*-CRP datasets (Fig. 2D, bottom plot).

Interestingly, epithelial maturation genes (e.g., cadherins, adherens junction proteins, and tight junction proteins), were both enriched and depleted in *hand2*-CRP datasets (Fig. 2D, middle plot). Of course, epithelial maturation is not specific to heart development; many derivatives of the LPM and IM undergo epithelial maturation as a part of their differentiation and morphogenesis (Drummond 2000, Trinh and Stainier 2004). For example, kidney progenitors aggregate and undergo epithelial maturation to generate the pronephros tubules in a *Wnt-4* dependent manner (Kispert, Vainio et al. 1998). We hypothesized that the enrichment of some epithelial maturation genes in the *hand2-CRP* datasets reflected their involvement in pronephros epithelial maturation. Consistent with this, the tight junction protein *tjp1b,* previously found to be excluded from renal progenitors, was depleted in the *hand2*-CRP datasets; *tjp2a*, which is enriched in the pronephros (Kiener, Sleptsova-Friedrich and Hunziker 2007), was enriched (Fig. 2D, middle plot). This pattern of enrichment and depletion suggests that Hand2-regulated epithelialization within the LPM is dependent on the progenitor cell-type: Hand2 activates cardiac progenitor epithelial maturation genes and represses pronephros progenitor epithelial maturation genes.

To identify biological processes associated with the positive and repressive roles of Hand2, we performed gene ontology (GO) enrichment analyses on depleted (downregulated) and enriched (upregulated) DEGs in *hand2-CRP* samples (Fig. 2E, Table S4). GO term analysis of downregulated DEGs (genes positively regulated by Hand2) revealed biological processes related to cardiac development including “atrial cardiac muscle tissue development”, “branching in blood vessel morphogenesis” and “sarcomere organization” (Fig. 2E, top plot, blue bars). In contrast, upregulated DEGs (genes negatively regulated by Hand2) showed enrichment in biological processes associated with ectodermal and IM derived tissues, including “regulation of anterior head development”, “neurocranium formation”, “otic placode formation, and “kidney development” (Fig. 2E, bottom plot, red bars).

To further explore the dichotomy of enriched GO terms for upregulated and downregulated DEGs, we categorized GO terms based on the tissue in which they normally function within the embryo (“LPM” vs “non-LPM”). For example, “atrial cardiac muscle tissue development” is a biological process that occurs in tissues derived from the LPM; “pronephric development” only occurs in IM-derived (non-LPM) tissues (Fig. 2E, Table S5). DEGs associated with epithelialization were classified as “epithelial-related” if they were associated with migration, cell polarity, or epithelial identity (Fig. 2E, Table S5, yellow bars). This classification scheme revealed that downregulated genes in *hand-CRP* datasets were non-specific, as they were enriched for GO terms associated with LPM, non-LPM, and epithelial-related biological processes; in contrast, the majority of upregulated genes in *hand-CRP* datasets were enriched for non-LPM GO terms (Fig. 2E, Table S5, red bars). This significant enrichment of GO terms further supports Hand2 functions as an important repressor of non-LPM biological processes.

### ATAC-seq identifies Hand2 dependent chromatin accessibility changes

To determine the mechanism by which gene expression is modulated by Hand2, we analyzed chromatin accessibility using ATAC-seq in LPM cells isolated from wildtype and *hand2*-CRP *Tg(hand2enh:eGFP)* embryos. We examined the tissue specificity of the ATAC-seq data by assessing the chromatin accessibility of LPM and non-LPM genes. As expected, genes known to be expressed in the LPM, such as *gata5* and *gata6*, contained ATAC-peaks in their promoters and exonic regions; genes not expressed in the LPM, such as *foxn1*, did not (Fig. 3A). Comparison of peak profiles, between wildtype and *hand2*-CRP samples, revealed a similarity between the datasets (Fig. 3B). ATAC peaks near promoter regions centered at transcription start sites (TSS) in both wildtype and *hand2*-CRP datasets and showed overlapping profiles (Fig. 3B). The similar distributions of fragment lengths for the wildtype and *hand2*-CRP libraries supports consistency between the datasets (Fig. S7). These characterizations of the ATAC data indicate that the wildtype and *hand2*-CRP datasets are of comparable high-quality data that can be used for differential analysis.

**Fig. 3:**
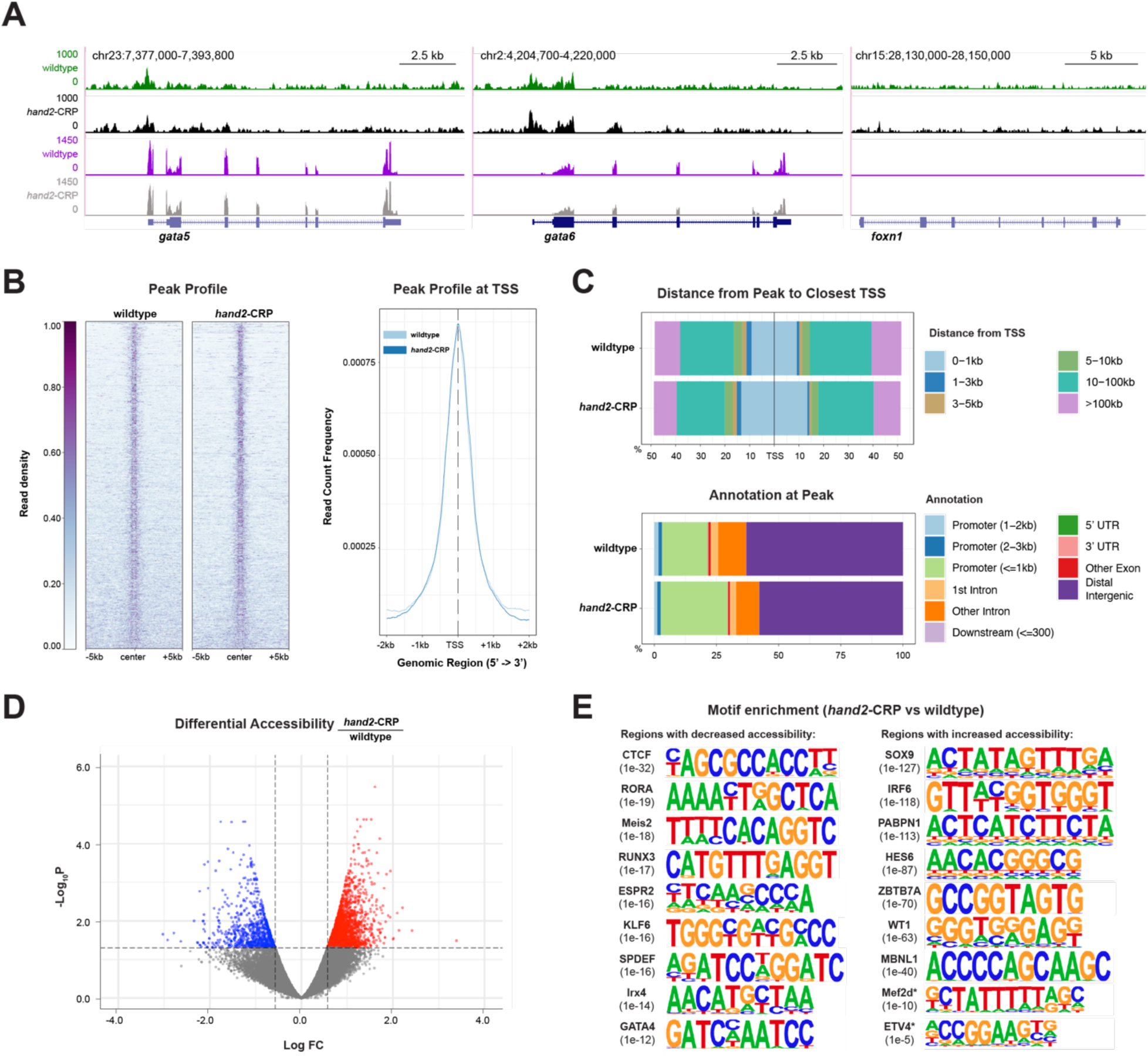
Genome-wide accessibility changes in *hand2*-CRP datasets compared to wildtype. (A) UCSC Genome Browser screenshot for *gata5*, *gata6*, and *foxn1* in wildtype and *hand2* crispant (*hand2*-CRP) LPM cells. Tracks are scaled to the same reads per kilobase per million (RPKM) values as indicated on the y-axis. The top two tracks are ATAC data (green, wildtype; black, *hand2*-CRP); the bottom two tracks are from RNA-seq (purple, wildtype; gray, *hand2*-CRP). (B) Heat map (left) and average profile (right) of ATAC fragments from wildtype and *hand2*-CRP LPM cells. Heat map (left) showing the read density 10kb around the peak centers of each fragment sorted and normalized by maximum RPKM. Average profile (right) of ATAC peaks from wildtype (light blue) and *hand2*-CRP (dark blue) samples occurring within 2kb of TSSs. (C) Distance from transcriptional start site (TSS) (top plots) and genomic annotation of ATAC peaks (bottom plots) in wildtype and *hand2*-CRP samples, characterized as a percentage of total peaks (15,033 wildtype peaks; 13,154 *hand2*-CRP peaks). (D) Volcano plot of relative accessibility level of significantly differentially accessible chromatin regions (DARs) in *hand2* crispant relative to wildtype samples (p-val < 0.05, log_2_FC ≥ |0.5|, dotted line, red dots (enriched in *hand2-CRP*) or blue dots (depleted in *hand2-CRP*)). Each dot corresponds to an individual chromatin region, with gray dots indicating nonsignificant DARs. (E) *De novo* motif enrichment of ATCT peaks with decreased accessibility (-DARs, left) and increased accessibility (+DARs, right) in *hand2-CRP* samples. Selected enriched motifs are shown next to the transcription factor with the most similar motif. P-value (in brackets) indicates significance of enrichment relative to the background. The two motifs, Mef2d and ETV4, that perfectly match known transcription factor binding sites are indicated by a *.

Gene expression changes can be regulated through changes in chromatin accessibility at promoters and distal cis-regulatory elements (Andersson 2015). To determine whether Hand2 modulates chromatin accessibility in specific regions of the genome, we examined the ATAC peak distribution in wildtype and *hand2*-CRP datasets across the genome. We mapped the ATAC peaks based on genome annotation and distance from the closest TSS (Fig. 3C, Table S6). The *hand2*-CRP datasets had a greater proportion of ATAC peaks within 1kb of a TSS (26.8%,) compared to wildtype datasets (18.4%; Fig. 3C, top). A similar enrichment of ATAC peaks in *hand2*-CRP (26.9%) was observed when comparing peaks within 1kb from annotated promoters (Fig. 3C, bottom). Consistent with this shift, wildtype datasets showed a slightly larger proportion of distal peaks (more than 10kb away from a TSS) than observed in the *hand2*-CRP datasets (69.1 % vs 62.1%); wildtype dataset showed a larger percentage (63%) of ATAC peaks in distal intergenic regions of when compared to *hand2*-CRP datasets (63% vs 57.7%). This shift in chromatin accessibility in the absence of Hand2 function, from distal intergenic regions to regions more proximal to promoters, suggests that Hand2 plays its repressive role through altering chromatin accessibility at distal regulatory elements.

While the genomic distribution of ATAC peaks reveal differences in the portion of accessible regions in *hand-*CRP and wildtype datasets, such analyses do not allow direct comparisons of individual peaks between the two datasets. A differential accessibility analysis between wildtype and *hand2*-CRP datasets identified 3,349 significantly differentially accessible regions (DARs, Fig. 3D, Table S7). Of these, 797 regions decreased in accessibility (-DARs) in *hand2*-CRP samples, while 2,552 regions increased in accessibility (+DARs). This preponderance of genomic regions with increased accessibility in the absence of Hand2 function is consistent with Hand2 directing repressive chromatin remodeling in wildtype LPM cells.

Chromatin accessibility changes are typically directed by transcription factor interactions with co-activators, repressors, or chromatin modulators (Long, Prescott et al. 2016). Therefore, we performed *de novo* motif enrichment analysis on DARs to identify potential binding sites for known transcription factors (Fig. 3E, Supplemental Data). Motif enrichment analysis on the 797 -DARs identified binding site motifs for transcription factors known to either interact with Hand2 and/or to regulate cardiac development (Fig. 3E, left column): Irx4, Gata4, retinoic acid-related orphan receptor-α (Rora), Meis2, and Runx3. Hand2 directly regulates *Irx4* and physically interacts with Gata4 to control cardiac gene expression (Bruneau, Bao et al. 2000, Dai, Cserjesi et al. 2002); Rora and Meis2, have been implicated in cardiac development and morphogenesis (Machon, Masek et al. 2015, Hill, Demarest et al. 2017, Beak, Kang et al. 2019, Nakajima 2019). Hand2 is thought to play a role in the repression of osteoblast differentiation by Runx3 and Runx2 (Yoshida, Yamamoto et al. 2004, Funato, Chapman et al. 2009). In addition, we found significant enrichment of binding sites for the insulator protein, CTCF, in -DARs (Fig. 3E, Supplemental Data). Enrichment of the CTCF motif is consistent with -DARs containing enhancers, as CTCF can facilitate enhancer-promoter interactions via chromosome looping (Nora, Goloborodko et al. 2017, Ren, Jin et al. 2017).

Motif enrichment analysis on +DARs identified binding sites for transcription factors implicated in biological processes associated with Hand2-repressed genes and with non-cardiac tissue development (Fig. 3E, right column). GO enrichment analyses identified head and kidney development as biological processes among +DARs, in addition to transcription factors involved in neurocranium morphogenesis (Irf6 and Hes6) and skeletal development (Sox9, Pabpn1, Mbnl1, Mef2d) (Kreidberg, Sariola et al. 1993, Kanadia, Johnstone et al. 2003, Michel-Olivier, Elena et al. 2003, Carroll, Macias Trevino et al. 2020). Among enriched transcription factor binding motifs in +DARs was the binding motif for ETV4, an ETS-domain family transcription factor that indirectly interacts with Hand2 in limb development through physical interaction with TWIST1 (Fig. 3E) (Firulli, Krawchuk et al. 2005, Zhang, Sui et al. 2010). Taken together, the analysis of -DARs and +DARs suggests that Hand2 acts with co-factors to modulate chromatin accessibility, promoting cardiac development and repressing non-cardiac developmental processes within the LPM.

To gain insights into the relationship between Hand2-dependent chromatin remodeling and gene expression, we assessed whether differentially expressed genes (DEGs) reflect chromatin remodeling events. Despite Hand2 samples having a larger proportion of ATAC peaks mapping to promoters and TSS (Fig. 3C), we found that chromatin accessibility around DEG promoters remained relatively unchanged, with only 2.3% of depleted genes and 0.84% of enriched genes exhibiting significant accessibility changes within 2kb of their TSS (Fig. S8). These data suggest that Hand2 is not regulating DEG expression through chromatin remodeling of their promoters or TSS.

### Identification of potential Hand2 dependent enhancers

The physical size and prediction of gene-enhancer interactions point to the DARs as potential enhancers. DAR lengths are within the range of functional enhancers. Enhancers are typically 50 - 1,500 bps, increasing in length with the number of transcription factor binding sites (Blackwood and Kadonaga 1998). 73% of our DARs were between 300 - 1,500 bps, while 93% were between 300 - 3,000 bps (Fig. 4A, Table S7), well within the expected enhancer size distribution.

**Fig. 4:**
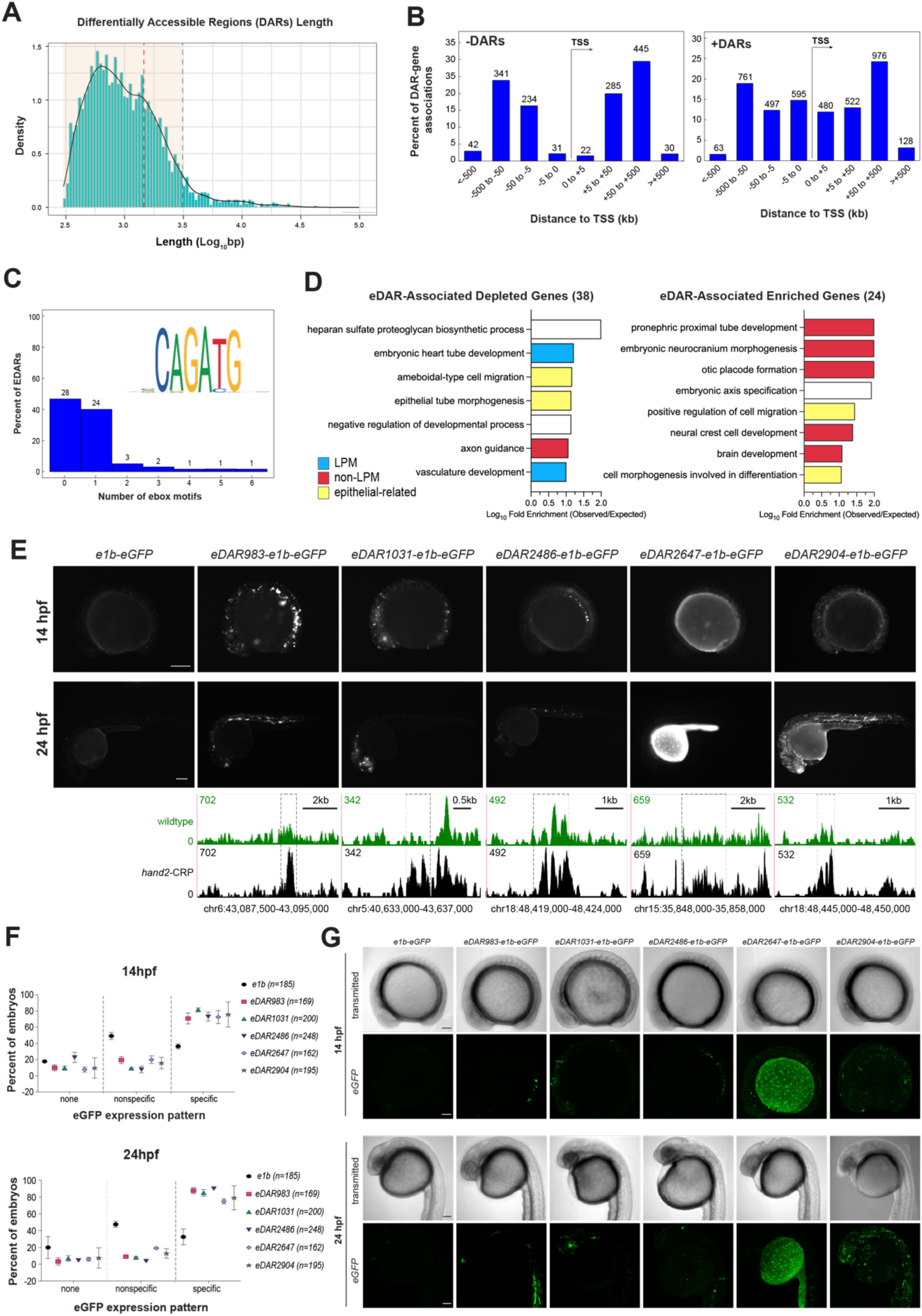
Differential Accessible Regions (DARs) contain functional enhancers. (A) Length distribution of significant DARs with 100bp bins. Orange window indicates expected size for an enhancer (300-3,000bp), with the upper size ranges expected for a low-complexity enhancer (red dotted line; 1,500bp) or a high-complexity enhancer (black dotted line; 3,000bp) indicated. (B) Position of DARs relative to the TSS of associated genes as determined by GREAT analysis. More DARs with increase accessibility in *hand2-CRP* samples are positioned within +/- 5kb of the TSS. (C) *Ebox* motif enrichment analysis was performed on 60 DARs that associate with differentially expressed genes (eDARs). The Hand2 consensus sequence logo from JASPAR (MA1638.1) shown in the upper right-hand corner. (D) GO enrichment analysis for biological process of genes associated with eDARs: 38 genes depleted and 24 genes enriched in *hand2-*CRP samples. GO subclasses with Log_10_ fold enrichment > |1.0| were classified as either LPM (processes occurs only in LPM, cyan), non-LPM (processes excluded from the LPM, red), or epithelial-related (processes involved in regulation of epithelial cell polarity, yellow). General biological processes found in both LPM and non-LPM tissues are indicated in white. (E) Functional enhancer reporter assay with epifluorescence images for select eDARs containing at least one *Ebox* motif. Embryos exhibiting specific expression patterns for each condition imaged at 14hpf (developmental stage when eDARs were identified, upper panels) and 24hpf (developmental stage with distinguishable organ primordia, lower panels) with the same exposure time across all samples. UCSC Genome Browser tracks for wildtype (green) and *hand2*-CRP (black) ATAC data at the endogenous eDAR loci are shown under the corresponding reporter images. Region corresponding to eDARs are indicated by a dotted box. Tracks are scaled to the highest RPKM value at each locus (top left corner). (F) Quantification of *eGFP* expression patterns in embryos injected with indicated enhancer reporter constructs (N = 3). Total number of embryos for each eDAR indicated in legend to the right (*n*). (G) Confocal maximum intensity projection images of eDAR-injected embryos with specific expression patterns. Scale bars are 100um in E and G.

Genomic Regions Enrichment of Annotations Tool (GREAT) allowed us to identify potential gene-enhancer relationships from the DARs (McLean, Bristor et al. 2010). GREAT has been successfully used to identify both short-range and long-range functional relationships between enhancers and their target genes (McLean, Bristor et al. 2010). GREAT predicts functional relationships between *cis* regulatory elements and the genes they may regulate by assigning a “regulatory domain” for every gene. Regulatory domains can be broken down into “basal” domains, which are defined as between 1kb downstream and 5kb upstream from a gene’s TSS, and “extended” domains, which can be positioned up to 1Mb upstream or downstream from a gene’s TSS. Chromatin regions that fall within a gene’s assigned regulatory domain can be flagged as a potential gene-enhancer interaction.

GREAT analysis with the 3,198 DARs identified a total of 5,452 putative DAR-gene interactions. The majority of DARs were predicted to interact with at least 2 genes (Fig. 4B), as seen in other enhancers (Nickol and Felsenfeld 1988, Grossniklaus, Pearson et al. 1992, Gómez-Skarmeta, Rodríguez et al. 1995, Spitz, Gonzalez et al. 2003, Cajiao, Zhang et al. 2004). Interestingly, when examining the distance between DARs and their putative target genes, we found that 26.7% of the gene interactions with +DARs occurred within 10kb of the gene’s TSS, far greater than the 3.71% of gene interactions with -DARs (Fig. 4B, Table S8). This is consistent with Hand2 negatively regulates chromatin accessibility at promoters (Fig. 3C, Table S6); however, these promoters do not appear to be associated with DEGs at the developmental stage when RNAseq was performed (Fig. S8).

ATAC-seq data have been used across a wide variety of organisms to identify *cis* regulatory elements that contain transcription factor binding sites (Daugherty, Yeo et al. 2017, Quillien, Abdalla et al. 2017, Yuan, Song et al. 2018, Bozek, Cortini et al. 2019). Despite strongly correlating with active enhancers and promoters, accessibility data do not necessarily inform gene expression level as chromatin accessibility changes can precede changes in transcription of target genes (Maybury-Lewis, Brown et al. 2021, Lin, Swedlund et al. 2022, Xiong, Tolen et al. 2022). To identify candidate enhancers that may be involved in Hand2-dependent gene expression changes at the early stage of CPC differentiation, we focused our efforts on the GREAT-predicted gene interactions for DARs associating with differentially expressed genes. From the DAR-gene associations predicted by GREAT, we identified 60 DARs associated with 62 genes showing differential expression between wildtype and *hand2-*CRP samples at the stages when the RNAseq was performed (14hpf; Table S8). We classified these as potential enhancer DARs (eDARs) as they were not located within promoter regions. Motif analysis of the eDARs identified 32 eDARs containing at least one *ebox*, the Hand2 consensus binding site (Fig. 4C) (Dai and Cserjesi 2002), suggesting that Hand2 may directly interact with these regions to regulate chromatin accessibility.

We performed GO term enrichment on the differentially expressed genes associated with eDARs to determine what biological processes are potentially regulated by eDARs (Fig. 4D). As expected, the eDAR-associated genes depleted in *hand2*-CRP samples showed an enrichment for LPM-derived biological processes such as “embryonic heart tube” and “vasculature development” (Fig. 4e, left panel). Additionally, enriched GO terms included biological processes related to formation of the myocardial epithelium (Fig. 4D, left panel). Conversely, eDAR-associated genes upregulated in *hand2*-CRP samples were enriched for non-LPM biological processes including “pronephric proximal tube development,” and “embryonic neurocranium morphogenesis” (Fig. 4D, right panel). Thus, GO-term analysis of the DEGs associated with eDARs suggest that Hand2 may have a role in positively regulating genes involved in LPM and negatively regulating non-LPM processes through chromatin remodeling of associated eDARs.

### Functional testing of potential Hand2 dependent enhancers

GREAT analysis allowed us to identify potential enhancer-gene relationships between a select group of DARs and differentially expressed genes. While Hand2’s role as an activator of myocardial cell fate is well established, its role as a repressor of the kidney lineage has only been recently described (Perens, Garavito-Aguilar et al. 2016). DEGs enriched in *hand2-CRP* samples associated with eDAR are significantly enriched for non-LPM biological processes, including pronephros development (Fig. 4D, right panel). We therefore sought to further explore Hand2’s repressive role in non-LPM processes.

We performed enhancer reporter assays in zebrafish embryos to verify GREAT-predicted +eDARs containing *ebox* motif function as enhancers (Fig. 4E). We injected zebrafish embryos with constructs that incorporated eDARs associated with genes enriched in *hand2*-CRP samples into an *eGFP* reporter construct containing the e1b minimal promoter. Enhancer activity was scored by examining the *eGFP* expression level at early (14 hpf) and late (24 hpf) stages. The early stage corresponds to the developmental stage from which eDARs were identified, and the expected onset of reporter expression. By the late stage, embryos have developed distinguishable organ primordia including pronephros tubules, brain divisions, heart, retina, and olfactory placode. Scoring this late stage allowed us to determine the primary tissue of *eGFP* expression. We found that *eGFP* expression increased substantially in eDAR-injected embryos compared to those injected with only the minimal e1b promoter as determined by fluorescence imaging (Fig. 4E). The increase in expression was observed at both 14 hpf and 24 hpf, with reproducible expression patterns across individual embryos and experiments (Fig. 4E-G).

Each eDAR drove unique tissue-specific reporter expression patterns in injected embryos (Fig. 4F). All +eDAR reporter constructs exhibit expression in tissues outside of the LPM. The basal promoter, *e1b*, exhibited a low level of basal expression in the notochord and somites (36% of *e1b-eGFP* injected embryos) and nonspecific expression patterns that varied significantly from embryo to embryo (49% of *e1b-eGFP* injected embryos) (Fig. 4E-G). In contrast, *eDAR983-e1b-eGFP* injected embryos had strong expression in the paraxial mesoderm at 14 hpf (71%) and somite-derived muscles and olfactory placode at 24 hpf (88%). Embryos injected with *eDAR1031-e1b-eGFP* had reporter expression in the forebrain, midbrain, and retina at both 14 hpf (82%) and 24 hpf (85%). The IM (73%) and, later, pronephric tubules (90%) were strongly labeled in *eDAR2486-e1b-eGFP* embryos while *eDAR2647-e1b-eGFP* embryos had reporter expression in the yolk syncytial layer (73% at 14 hpf, 75% at 24 hpf). Finally, the vast majority (76% at 14 hpf, 79% at 18 hpf) of *eDAR2904-e1b-eGFP* injected embryos exhibited reporter expression in the majority of tissues with the exception of the LPM. Robust *eGFP* expression in the LPM was not observed in embryos injected with +eDAR reporters when screened by epifluorescence microscopy. Together, these findings demonstrate that Hand2-dependent chromatin regions are functional enhancers; however, these enhancers do not drive detectable expression in the LPM, a tissue from which the eDARs were identified.

The lack of robust reporter expression in the LPM and broad reporter expression pattern of the +eDARs tested led us to hypothesized that Hand2 expression in the LPM may prevent enhancer activation for the +eDARs within this tissue. To test whether Hand2 was required for the +eDAR repression, we injected *eDAR1031-e1b-eGFP* into *hanS6* embryos, which lack Hand2 function, and evaluated the LPM for *eGFP* expression (Fig. 5A). We found that the +eDAR reporter constructs were significantly more likely to express *eGFP* in the LPM of *hanS6* embryos then the LPM of wildtype siblings (Fig. 5B). Taken together, these data demonstrate that Hand2 is sufficient to repress these enhancers within the LPM.

**Fig. 5:**
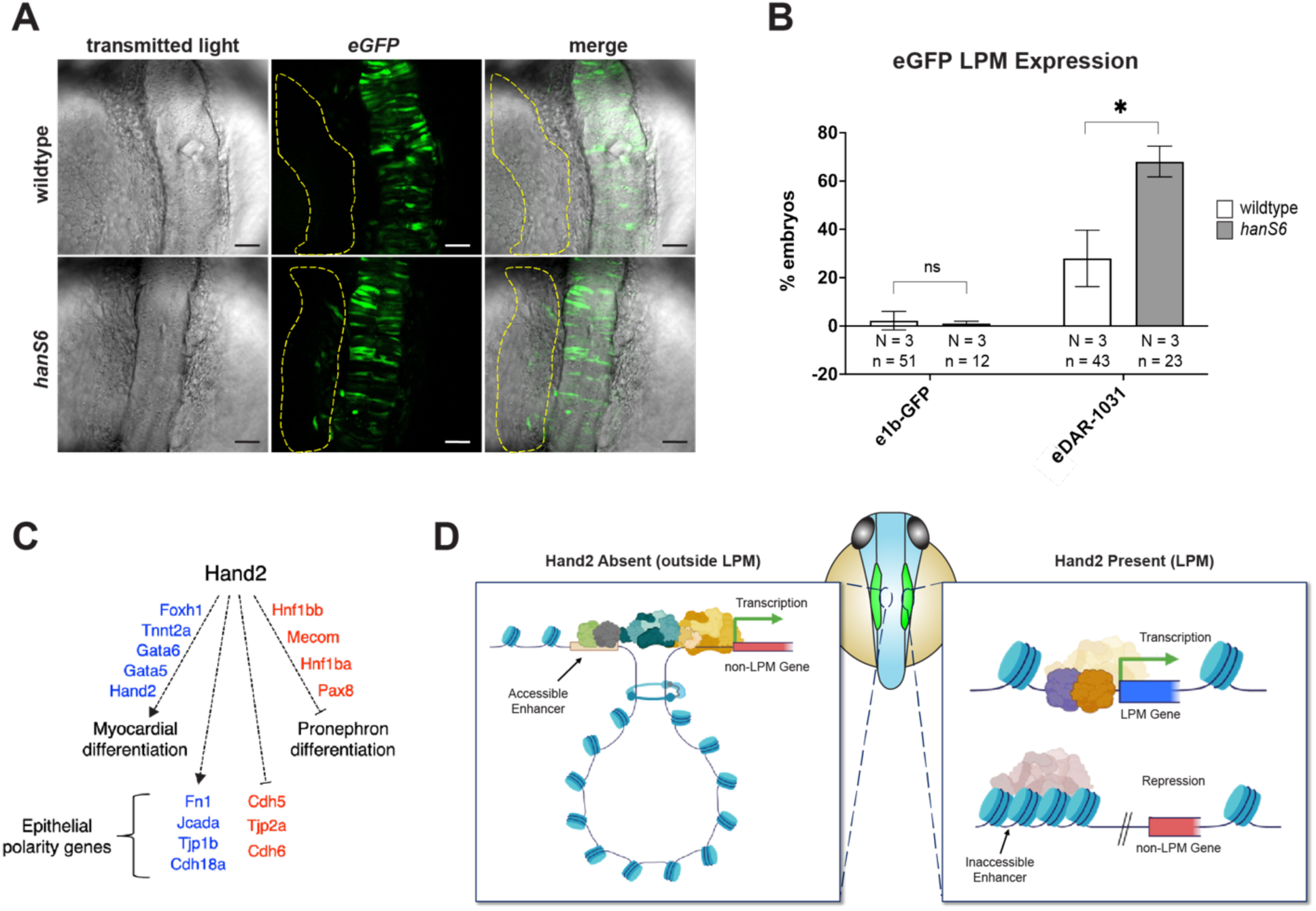
Hand2 represses enhancer activity of differential accessible regions. (A) Confocal images of wildtype and *hanS6* 14 hpf embryos injected with the *eDAR1031-e1b-eGFP* reporter construct, showing reporter gene expression in the LPM (yellow dashed region) only in the *hand2* mutant embryo (lower panel). Scale bars are 50um. (B) Quantification of wildtype and *hanS6* embryos injected with *eDAR1031-e1b-eGFP* that exhibit *eGFP* expression within the LPM. “N” denotes the number of independent experimental replicate. “n” denotes the total number of embryos quantified. *eGFP* expression from embryos injected with the *e1b-eGFP* construct are shown as a control (*, p-value < 0.05, ns = not significant). (C) Putative Hand2-regulated gene network compiled based on the analysis of genes that are downregulated (blue text) or upregulated (red text) in the absence of Hand2. Lines with arrowheads signify an activator function for Hand2 in the indicated biological process while lines with a block arrow signify a repressive function for Hand2 in the indicated biological process. Dashed lines indicate Hand2 may either directly or indirectly regulate list genes. (D) Proposed mechanism of Hand2-dependent regulation of non-LPM and LPM-derived tissues based on the results of this study. Schematic of a 14 hpf zebrafish embryo is shown in the middle with the LPM colored in green. The embryo proper is colored blue while the yolk is colored beige. In the LPM, Hand2 positively regulate expression of LPM genes and limit the accessibility of non-LPM genes. In non-LPM tissues where Hand2 is not expressed, enhancers of non-LPM genes are accessible and can act on target non-LPM genes.

## Discussion

By differential analyses of wildtype and Hand2-deficient LPM cells at the onset of cardiac differentiation, using RNA- and ATAC-seq, we identified both positive and negative roles for Hand2 in regulating cardiac and non-cardiac biological processes, respectively. Our analyses revealed genes involved in myocardial differentiation and epithelial maturation that were previously unknown to be Hand2-dependent, expanding the Hand2-regulated cardiac differentiation gene network (Fig. 5C). Additionally, we find a repressive role of Hand2 in pronephros differentiation, finding that Hand2 represses critical genes such as *hnf1bba*, *hnf1bb*, *mecom*, and *pax8* (Fig. 5C). Through combining differential chromatin accessibility analysis with gene expression, we identified functional enhancers whose chromatin accessibility is repressed in the presence of Hand2. The identified enhancers are predicted to associate with Hand2-dependent genes, suggesting that Hand2 represses non-LPM genes within the LPM through chromatin remodeling of these enhancers.

Based on the findings, we propose that Hand2 coordinates disparate developmental programs to ensure proper cardiac morphogenesis (Fig. 5D). Outside of the LPM where Hand2 is not expressed, its absence fosters open chromatin, and this accessibility permits enhancers to activate genes important for the development of paraxial mesoderm, IM, and chordamesoderm tissues (Fig. 5D, left panel). Within the LPM the presence of Hand2 fosters a closed chromatin conformation at these enhancers, reducing the potential activation of their associated genes (Fig. 5D, right panel).

Previous studies have shown that epithelial maturation is disrupted in Hand2 mutants (Trinh, Yelon et al. 2005, Garavito-Aguilar, Riley et al. 2010). Our transcriptome data in Hand2 mutant cells point to the epithelial maturation defect in cardiac progenitors at the transcriptional level. Furthermore, differential transcriptome analysis suggest that the epithelial gene expression program may be inherent to the development of a particular cell-type as we find that epithelial genes associated with myocardial progenitors are decreased while epithelial genes expressed in pronephric progenitors are increased in the absence of Hand2.

Our data indicate that Hand2 restricts fates within the LPM through chromatin accessibility of non-LPM enhancers. The Hand2-dependent accessible regions that function as enhancers are associated with genes known to be involved in biological processes that occur outside the LPM and drive reporter expression in non-LPM derived tissues. The majority of the biological processes identified by GO enrichment involve tissues that arise from LPM-adjacent cells, including the paraxial mesoderm (cartilage, myotome, and head structures), IM (kidney), and chordamesoderm (central nervous system). RNA-seq data from purified LPM cells show that genes associated with these non-LPM derived tissues including *lhx1a*, *vcana*, *pax2a*, *foxj1a*, and *rbm47* are expressed in the LPM at low levels. However, their expression increases in the absence of Hand2 indicating that Hand2 functions to dampen non-LPM gene expression within the LPM. Differential ATAC-seq indicates that Hand2 is generally repressive for chromatin accessibility (Fig. 3D). We found evidence for Hand2-repression of key genes implicated in the early morphogenesis of non-LPM processes. Reporter assays of Hand2-repressed chromatin revealed activity in the CNS, non-cardiac muscles, and renal system, suggesting that Hand2 may act on these enhancers to repress these cell fates in the LPM.

Our data suggest that Hand2-dependent repression of non-LPM genes through chromatin remodeling of their enhancers may occur through interactions with co-factors. Motif enrichment of ATAC-peaks with increase accessibility in the absence of Hand2 (Fig. 3C) identified binding sites for factors that regulate these pathways (Sox9, Wt1, Ebf, Hes6, Lin28a, Irf6) in Hand2-repressed chromatin regions. Hand2 contains both a basic DNA-binding motif and a dimerization motif, each of which play different roles during development (Schindler, Garske et al. 2014). In mice, Hand2’s dimerization domain is most critical for CPC differentiation while its DNA-binding ability plays a less central role (Liu, Barbosa et al. 2009). Hand2 forms a complex with Gata4 through dimerization with Twist1. It is Hand2 dimerization, and not direct DNA-binding, that activates gene expression (Schindler, Garske et al. 2014). This may explain why the Hand2 consensus motif was not enriched in most of the Hand2-dependent chromatin regions in either this or previous studies (Dai, Cserjesi et al. 2002, Firulli, Krawchuk et al. 2005). It is therefore likely that Hand2-dependent chromatin regulation during CPC differentiation is achieved through interactions with chromatin remodeling cofactors.

In addition to interaction with co-factors to modulate enhancer activity, Hand2 may repress a subset of non-LPM enhancers through direct DNA-binding. Several of the Hand2-repressed genes found in this study were previously shown to be associated with genomic regions directly bound by Hand2 in the limb bud of mice. These include *pax8*, *pax2*, *lhx1*, *cdh17*, and *mecom* (kidney), *foxj1*, *emx2*, and *crabp2* (nervous system), and *prdm1* (neurocranium morphogenesis) (Osterwalder, Speziale et al. 2014). We also found binding sites for SWI/SNF chromatin remodelers ZBTB7 and PRDM9 in Hand2-repressed chromatin regions (Koh-Stenta, Joy et al. 2014, Gupta, Singh et al. 2020). Recent work found that Hand2 DNA binding recruits PHF7, a member of the SWI/SNF chromatin remodeling complex, to cardiac enhancers (Garry, Bezprozvannaya et al. 2021). Finally, a subset of eDARs contain *Ebox* motifs, further supporting the hypothesis of direct Hand2-DNA binding to enhancers.

## Methods

### Zebrafish husbandry

Zebrafish were raised and maintained in accordance with recommendations from the Guide for the Care and Use of Laboratory Animals by the University of Southern California as described previously (Westerfield 2000). Protocol was approved by the Institutional Animal Care and Use Committee (IACUC). *Tg(hand2enh:eGFP)* and *Tg(myl7:nucGFP)* fish were outcrossed for all experiments. Wildtype embryos were obtained via incrosses. Embryos were raised in egg water (60mg/ml of Instant Ocean and 75mg/ml of CaSO_4_ in Milli-Q water) at 28.5°C. Larvae and fish older than 16-somites were anesthetized in 0.01% MS222, pH 7.2.

### Generation of transgenic lines

Upstream *cis*-regulatory elements were amplified from wildtype zebrafish genomic DNA and cloned into the e1b-eGFP-Tol2 vector (available from Addgene, Plasmid #37845) by restriction cloning at the XhoI and BglII sites. Primers used to amplify genomic regions are listed in Table S1. 2.5nL of pre-mixed constructs and *tol2* mRNA were co-injected each at a final concentration of 50ng/ul into one-cell stage zebrafish embryos. Injected F0s were raised to reproductive age, outcrossed with wildtype, and screened for expression. Founders with successful transposition events were identified using splinkerette nested PCR (Uren, Mikkers et al. 2009) and raised to adulthood. Lines used in this study are listed in Table S2.

### Enhancer reporter assay

Enhancer reporter assays were performed by injection into wildtype embryos at the one-cell stage with 2.5nL injection solution (25ng/ul reporter construct, 25ng/ul *tol2* mRNA, 10% phenol red (5mg/mL in 5mM KCl)). Embryos with gross morphological defects prior to 10 hpf were not included in expression evaluation. Expression patterns were evaluated at 14 hpf and 24 hpf and embryos imaged on a Zeiss Axio Zoom fluorescence stereo zoom microscope at 62X and 39.5X magnification and 300ms exposure time. Expression patterns were quantified by measuring the percentage of morphologically wildtype injected embryos with one of three expression patterns (none, nonspecific, or specific patterns). Statistical significance was determined by binning 50 - 70 embryos from different experimental days into a single “replicate” for a total of 3 replicates, then performing a student’s t-test (p > 0.05). Representative embryos were imaged on a Zeiss LSM 880 inverted microscope by embedding in 1% UltraPure Low Melting Point Agarose (Thermo Fisher, Catalog #16520050) in 1X Embryo media (17.4 mM NaCl, 0.21 mM KCl, 0.12 mM MgSO_4_, 0.18 mM Ca(NO_3_)_2_, 1.5mM HEPES pH 7.6). *eGFP* images captured using epifuorescence microscopy are pseudocolored white while confocal acquired images are green.

### CRISPR-Cas9 mutagenesis and morpholino knockdown

Target-specific Alt-R® CRISPR modified crRNAs and universal tracrRNAs were synthesized by Integrated DNA Technologies (Alt-R™ CRISPR crRNAs and tracrRNA, Integrated DNA Technologies, Coralville, IA, USA) and resuspended in IDT duplex buffer at 100uM stock solutions. Equivolume crRNA and tracrRNA 100uM stocks were mixed and annealed in a thermocycler using the following conditions (95°C 5min, cool to 25°C at 0.1°C/sec, 25°C for 5min) to form crRNA:tracrRNA duplexes. 250 ug of Cas9-NLS (PNABio, Catalog #CP02) was resuspended in 50ul nuclease-free water to form 30.68uM stock solution. dgRNPs were prepared for zebrafish embryo injections by mixing 0.5 ul crRNA:tracrRNA duplex with 0.5ul nuclease-free duplex buffer (IDT catalog #11-0501), 0.81ul Cas9-NLS, and 2.69ul nuclease-free water to form 5uM dgRNP complexes. dgRNPs were incubated at -20°C overnight before use. On the day of injection, dgRNPs were heated to 37°C for 5 minutes and cooled on ice before adding 0.5ul 5mg/ml phenol red solution in 5mM KCl. 2.6nl was injected into one-cell stage embryos. Morpholino oligo against *hand2* (HAND2-MO, 5’-CCTCCAACTAAACTCATGGCGACAG-3’) was ordered from Genetools and resuspended in 5 mM HEPES pH 7.6. Embryos were injected with 2.5nl of either 50uM, 100uM, or 150uM morpholino at the one-cell stage.

### Capillary electrophoresis and fragment analysis

Genomic DNA was extracted by submerging individual 5 dpf anesthetized embryos in 20ul of 50mM NaOH solution and heating at 95°C for 10 minutes. Embryos were then cooled to 4°C for 10 minutes and 2ul of 1M Tris-HCl pH 8.0 was added to neutralize the solution. 1ul of genomic DNA solution was used in subsequent PCR reactions. CRISPR-Cas9 target sites were amplified from the genomic DNA of each individual embryo, with each forward primer labeled with 6-carboxyfluorescein (6-FAM, IDT) at the 5’ end. LA Taq® high fidelity DNA polymerase (TaKaRa, Catalog #RR002A) was used to amplify the CRISPR-Cas9 target site according to the manufacturer’s instructions. Amplicons were diluted to 0.5ng/ul and run-on ABI 3730xl sequencers by Azenta Life Sciences using the ABI DS333 dye set, which utilizes the GeneScan™ 500/1200 LIZ™ dye size standard. Data were analyzed using Peak Scanner CE Fragment Sizing software (Thermo Fisher), which scales fluorescent intensities from 0 to 33,000. Off-scale peaks, peaks appearing before 40 base pairs, and peaks with a combined RFU under 1,900 were considered background and discarded. The proportion of target sites with indel mutations was calculated as described previously (Hoshijima, Jurynec et al. 2019).

### Immunolabeling and Imaging

Live embryos were immobilized with 0.02% tricaine (Sigma, Catalog #A5040) and embedded UltraPure 1% low melt agarose (Thermo Fisher, Catalog #16520100) in 1X embryo media in PELCO Glass Bottom Petri Dishes (Ted Pella, Catalog #14027) in) and imaged on a Zeiss LSM 880 inverted confocal microscope. Embryos prepared for immunostaining were fixed overnight at 4°C in 2% PFA (Electron Microscopy Sciences, Catalog #15710). Embryos were either embedded in 4% low-melt agarose/PBS + 0.1% Tween and cut into 200um sections using a Leica VT1200 or stained as intact whole-mounts. Whole-mounts or sections were stained in PBDT (1X PBS, 1% BSA, 1% DMSO, 0.1% Triton X-100, pH 7.3). Embryos were de-yolked in PBDT using forceps prior to sectioning. Mouse monoclonal IgG_2a_ anti-PKC ζ (Santa Cruz Biotechnology, sc-17781) was used at 1:1000 to visualize aPKCs localization. Sections and whole mount samples were imaged on an inverted Zeiss LSM 880. Whole mount *Tg(myl7:NLS-GFP)* samples were imaged using a Zeiss LSM 780 upright. Resolution, pixel dwell, and pinhole size were kept consistent across embryos while laser power was adjusted from 0.5 - 6%. Imaging data were analyzed using Imaris (Bitplane) and ImageJ (FIJI).

### Embryo dissociation and flow cytometry

200 *eGFP*-positive wildtype and *hand2*-CRP *Tg(hand2enh:eGFP)* embryos were dechorionated in egg water. Embryos were then dissociated at 14 hpf in 1ml of 10mg/ml freshly prepared protease solution (Protease from *Bacillus licheniformis* Sigma-Aldrich: P5380) in DMEM/F-12 (Gibco catalog: 11039021) by incubating at 4°C for 1hr. Dissociated embryos were transferred to an Eppendorf tube by filtering through a 40um filter, pelleted, and resuspended in FACS buffer (1x PBS, 2% FBS, 1mM EDTA). Cells were stained with 1:1000 of 1mg/ml DAPI to identify dead cells. Debris and doublets were removed by gating. Approximately 10,000 *eGFP*-positive, DAPI-negative cells for each condition were sorted using a BD FACS Aria III cell sorter into 1x PBS 10% FBS. A subset of these cells was re-sorted to confirm purity of sorted cells (Fig. S4). Sorted cells were split and prepped for either RNA- or ATAC-library preparation and sequencing.

### RNA-sequencing and analysis

RNA from approximately 20,000 FACS-sorted cells was extracted using the RNAqueous Total RNA Isolation Kit (Invitrogen catalog: AM1912). Strand-specific libraries were constructed using the SMARTer Stranded Total RNA-seq Kit (Takara catalog: 634862) following ribosomal RNA removal using RiboGone (Takara catalog: 634846). Sequencing was performed on an Illumina HiSeq 2500 with 40 million 75bp paired-end reads. Sequences were aligned to the zebrafish genome (GRCz11) using STAR (Dobin, Davis et al. 2013). Transcript abundance was quantified using RSEM (Li and Dewey 2011). Biological replicates of the wildtype and *hand2-CRP* samples are similar to each other. Scatter plot comparison of the complete datasets indicating the purification and library production approach are highly reproducible (Fig. S5). DESeq2 was used to identify differential expression, calculate statistical significance of differentially expressed genes, and perform data visualization (Love, Huber et al. 2014). DEG expression levels are quantified as transcripts per million (TPM) + 1 to permit log scaling for visualization.

Gene ontology enrichment was performed on query gene lists using the Panther Classification System (Mi, Muruganujan et al. 2013). Statistical overrepresentation tests were performed on ENSDARG identifiers with all *Danio rerio* genes as background. P-values were calculated using Fisher’s Exact test and adjusted for False Discovery Rate (FDR < 0.05). Significant GO terms were sorted hierarchically according to related classes within an ontology, with only the most specific subclass considered for further analysis. Fold enrichment of statistically significant GO terms (FDR < 0.05) were calculated by dividing the number of genes per term by the expected number of genes per term given the size of the input list. GO terms were manually assigned to either LPM specific, non-LPM specific, epithelial-related, or general biological processes based on the tissue within which a given GO term occurs. For example, *pdgfrb*, which is associated with GO terms “tube morphogenesis” and “tube development” was classified as “epithelial-related” while *atf2*, which is only associated with general terms such as “regulation of cellular process”, “regulation of biological process”, and “biological regulation”, would not be categorized. Full list of GO terms (Table S4) and assigned GO classifications (Table S5) can be found in the supplement.

### ATAC-sequencing and analysis

Approximately 20,000 FACS-sorted cells were lysed in cold lysis buffer (10 mM Tris-HCL, pH 7.4, 10 mM NaCl, 3 mM MgCl_2_, 0.1% IGEPAL CA-630). ATAC-seq libraries were generated as described previously (Buenrostro, Giresi et al. 2013) using the Nextera XT DNA Library preparation kit (Illumina catalog: FC-131-1024). Sequencing was performed on an Illumina HiSeq 2500 with 30 - 40 million 75bp paired-end reads. Adapter and transposon sequences were trimmed using TrimGalore (https://github.com/FelixKrueger/TrimGalore). Reads were mapped to GRCz11 using bowtie2 with a 100bp - 2kb fragment threshold (Langmead and Salzberg 2012). Reads mapping to mitochondrial DNA were removed using samtools (Li, Handsaker et al. 2009).

Subsampling based on estimated library complexity was performed using ATACseqQC (https://jianhong.github.io/ATACseqQC/articles/ATACseqQC.html). PCR duplicates were removed using Picard (http://broadinstitute.github.io/picard/). To correct for Tn5 insertion, reads on the positive strand were shifted by +4bp while reads on the negative strand were shifted -5bp via ATACseqQC (https://jianhong.github.io/ATACseqQC/articles/ATACseqQC.html). The bamPEFragmentSize function from deepTools was used to calculate the size distribution of mapped ATAC fragments with 1bp-wide bins and 15bp smoothing (https://deeptools.readthedocs.io/en/develop/index.html).

Peak calling was performed with MACS2 (Zhang, Liu et al. 2008). RPKM-normalized bigWig files were generated using the bamCoverage function from deepTools and visualized using the UCSC Genome Browser (Kent, Sugnet et al. 2002, Ramírez, Dündar et al. 2014). Characterization of ATAC peaks and fragments were performed using the ChIPseeker package (Yu, Wang et al. 2015). We followed the *csaw* workflow pipeline (Lun and Smyth 2016) as described (Reske, Wilson et al. 2020). Overlapping MACS2 peaks were normalized using trimmed mean of *M* values, which eliminates biases in ATAC library distribution by assuming that the top and bottom quantiles of 10kb-binned counts are due to technical, and not biological, differences (Robinson, McCarthy et al. 2010). Significant (FDR < 0.05) DA peaks identified using *edgeR* (Robinson, McCarthy, and Smyth 2010) were annotated with Hypergeometric Optimization of Motif EnRichment (HOMER) (Duttke et al. 2019). HOMER was also used to find significantly enriched *de novo* motifs relative to the zebrafish genome (Duttke, Chang et al. 2019). Packages ggplot2 and pheatmap were used to generate histograms and heatmaps in RStudio (https://ggplot2.tidyverse.org/, https://r-charts.com/correlation/pheatmap/). For ATAC peak genomic distribution, only peaks with overlap between biological replicates were analyzed.

### Functional annotation of differential ATAC regions

GREAT version 3.0.0 was used as previously described to identify possible functional interactions between DARs and genes (http://great.stanford.edu/great/public-3.0.0/html/) (McLean, Bristor et al. 2010). Bed files containing the chromosome number, start, end, and seqID for DARs were sequentially converted from GRCz11 to danRer7 using the UCSC LiftOver Genome Annotations tool (https://genome.ucsc.edu/cgi-bin/hgLiftOver). Some DARs (151 regions) could not be lifted over to older genome assemblies and were excluded from the GREAT analysis. Analyses in Fig. 4B were generated using GREAT. DARs interacting with DEGs were subset from GREAT output files using RStudio. All GREAT output files can be found in Table S8.

## Acknowledgement

We would like to thank Eun (Daniel) Koo, Masahiro Kitano and Simon Restrepo for helpful discussions. We thank Edwin Carranza Lopez and Grey Thomas for expert fish care. We gratefully acknowledge the Translational Imaging Center (University of Southern Califorinia) for access to microscope and image processing software used in this work. In addition, we thank the Flow Cytometry Facility (University of Southern California, Stem Cell) and the Molecular Genomics Core (University of Southern California, Norris Comprehensive Cancer Center) for their assistance in this work.

## Competing Interest

The authors declare no competing or financial interests.

## Funding

This work was supported by a NIH award, R01HL140472 to LAT. VK was supported by a NIH T32 Chemistry-Biology Interface Training Grant (5T32GM118289). BC and RR were funded by NIH award R35GM130376 to RR. LAT and SEF were supported by University of Southern California.

## Data availability

Sequence data generated in this study have been submitted to GEO (GSE243290). The authors declare that all data and its supplementary information files within the article are available from the corresponding authors upon reasonable request.

## Notes

### Competing Interest Statement

The authors have declared no competing interest.

